# The European Neolithic Expansion: A Model Revealing Intense Assortative Mating and Restricted Cultural Transmission

**DOI:** 10.1101/2024.04.29.591653

**Authors:** Troy M. LaPolice, Matthew P. Williams, Christian D. Huber

## Abstract

The Neolithic revolution initiated a pivotal change in human society, marking the shift from foraging to farming. The underlying mechanisms of agricultural expansion are debated, primarily between cultural diffusion (knowledge and practices transfer) and demic diffusion, or people migration and replacement. Ancient DNA analyses reveal significant ancestry changes during Europe’s Neolithic transition, suggesting primarily demic expansion. However, the presence of 10-15% hunter-gatherer ancestry in modern Europeans indicates cultural transmission and non-assortative mating were additional contributing factors. We integrate mathematical models, agent-based simulations, and ancient DNA analysis to dissect and quantify the roles of cultural diffusion and assortative mating in farming’s expansion. Our findings indicate limited cultural transmission and predominantly within-group mating. Additionally, we challenge the assumption that demic spread always leads to ancestry turnover. This underscores the need to reassess prehistoric cultural expansions and offers new insights into early agricultural society through the integration of ancient DNA with archaeological models.

## Introduction

The transition to agriculture was transformative for human history, with pronounced effects on economics^1–3^, population size^4,5^, ancestry^6,7^, language^8,9^, and societal structure^5,10^. Deemed the “Neolithic Revolution”^11,12^, this change in behavior and culture set the basis for modern society, facilitating the construction of permanent settlements and structures within complex social networks centered around a more stable food and resource supply^10,13^. The shift to farming has also been associated with reductions in population health due to nutritional deficiencies and increased disease transmission^13–16^. As a result of the vast impacts of this change in societal structure and its health-related consequences, researchers have long been interested in reconstructing how and why the transition to agriculture occurred.

It is believed that early European agriculture originated in southwest Asia^17–19^ and first expanded into Europe from northwestern Anatolia^17^. One of the first attempts to describe this spread came in 1971, when Ammerman and Cavalli-Sforza used radiocarbon-dated sites to calculate the speed of the farming expansion across Europe (the so-called “*front speed”*)^20^. Ammerman and Cavalli-Sforza determined an average expansion rate of approximately 1 km per year^20,21^ more recently corroborated by Pinhasi *et al*., (2005)^22^.

In addition to estimating the front speed, studies have also sought to understand the specific mechanism that drove this expansion. One mechanism is *demic diffusion*, first coined in 1971 by Ammerman and Cavalli-Sforza^20^. Demic diffusion describes an expansion of agriculture driven by farmers migrating into previously un-farmed territories, thereby introducing their cultural practices as well as genetic ancestry into these new areas^20,21,23–25^. Ammerman and Cavalli-Sforza argued that the rate of farming expansion in Europe is principally compatible with a primarily demic model of population growth and displacement, and suggested that this would result in a genetic cline of Neolithic ancestry in the resulting populations^20,21^.

Conversely, under *cultural diffusion*, first discussed and applied to the Neolithic Expansion by Edmonson in 1961^26^, hunter-gatherer (HG) groups learn agriculture from farmers and acquire the means necessary to practice this culture by living near farmers^20,21,23–26^. Within the cultural diffusion process, both *vertical* and *horizontal* modes of transmission have been considered^25^. Horizontal transmission refers to individuals learning behaviors from peers in other populations, whereas vertical transmission is the passing down of cultural practices from parents to offspring^25^. Notably, in a cultural model, farming ancestry does not expand to the degree it does in a demic model. This is because farming expands predominantly by learning or imitation, instead of farmers (and their genomes) moving into and overtaking new territories.

Aside from the perhaps unrealistic extremes of a fully demic or fully cultural model, the two models can be combined into a *demic-cultural* mode of transmission, i.e. where farmers migrate into new land but also interact with HGs as they expand, either actively or passively passing knowledge onto those they meet^23–25^. Under this model, both mechanisms are thought to contribute, in varying degrees, to the expansion of farming.

Several previous studies have aimed to quantify the relative contribution of cultural vs. demic transmission under a combined demic-cultural model^23–25^. One measurement used to quantify the cultural contribution is “*cultural effect”*. Cultural effect is defined as the additional contribution of cultural transmission to the front speed on top of an otherwise demic mode of expansion, i.e. the relative increase in front speed due to cultural transmission^23–25^. Previous estimations have suggested that cultural effect could account for up to 40% of the spread of farming^23^, with others showing a maximum cultural effect of 21% to be consistent with the observed front speed^25^. In addition to the large uncertainty, these estimates do not consider patterns of genetic ancestry resulting from the respective models. This is important because a purely demic model (zero learning and full assortative mating) predicts a complete genetic turnover whereas cultural transmission allows for the persistence of indigenous HG ancestry after the expansion.

In addition to mathematical modeling of the Neolithic expansion^23–25,27–31^, other studies have gathered novel insights from investigating settlements^32–36^, geographical factors^37^, language^8^, ancient climate conditions^38^, and flora data^39^. In particular, recent analyses of ancient DNA (aDNA) have fundamentally transformed our understanding of population movements and associated genetic changes that took place during this period. With the advent of aDNA, researchers were able to scan for genetic adaptations in Neolithic populations^7^, determine distinct ancestry patterns and population movements^40^, or describe contributions of ancient groups to modern genomes^41^. Early work based only on mitochondrial data lent more support for a significant local Paleolithic HG contribution to modern European populations^42^ thereby indicating a primarily cultural transmission. However, ever-growing sample sizes, and especially whole-genome sequences, have revealed in greater detail a significant temporal and geographical spread of Anatolian farming ancestry across Europe^6,7,17,39,41^. But despite aDNA technology providing unprecedented resolution of genetic ancestry and population movements over time, a limitation of DNA is that it does not directly describe or indicate behavior, culture, or specific mechanisms of expansion. As such, an interdisciplinary approach linking behavioral mechanisms with genetic data is required to fully understand the Neolithic farming expansion.

Here, we provide such an approach through a demographic mathematical model and a spatial, agent-based, population genetic simulation. We model mortality under density-dependent competition based on carrying capacity estimates from literature^21,43^ and osteological age at death data from Neolithic samples^34^. We fit parameters in our model to archeological front speed estimates^20–22^ and ancestry estimates derived from 911 early European farming individuals. From this, we infer model parameters by determining the effect of dispersal rates on the front speed, and by tracing the genomic ancestry of each individual in our simulation. This allows us to elucidate which cultural transmission characteristics are compatible with archaeological and genetic data. We conclude that there must have been near complete cultural assortative mating within farming and hunter-gathering groups and very limited involvement of farmers in cultural transmission, with less than 0.1% of farmers converting an HG to farming annually. More generally, our modeling cautions that primarily demic expansion does not necessarily lead to a major ancestry turnover, suggesting that a naive interpretation of ancient ancestry patterns does not always reflect underlying behavioral mechanisms.

## Results

### Basic model suggests low levels of cultural transmission

We started by examining the influence of cultural transmission on the dispersal of Anatolian ancestry throughout a Europe inhabited by hunter-gatherers (HGs). We employed a reaction-diffusion model in a one-dimensional (1D) continuum, as a simplified representation of this process. This involves a system of partial differential equations that mirror those utilized in prior models addressing the spread of agriculture (e.g., Fort *et al*., 2012)^23^. Importantly, our model extends beyond these by incorporating the genetic ancestry of individuals at a single genetic locus, enabling us to trace the propagation of ancestral change over space.

We defined three population groups within the model: cultural farmers with Anatolian ancestry, cultural farmers with HG ancestry, and cultural HGs of HG ancestry. As an initial condition, we assumed that the farmers of Anatolian descent are constrained to 200 km at the left end of the range, and the HG group is constrained to the remaining 2,800 km of the range, with twenty times lower population density^21^. The spatiotemporal distribution of these groups is then influenced by three principal dynamics: random migration, which is determined by a diffusion constant; logistic population growth, modulated by growth rates and carrying capacities; and cultural transmission through learning. Our cultural transmission mechanism is informed by established theory^23^, considering a learning rate (*f*; the product of the number of teachers times the probability of learning per contact), and a bias parameter (γ), which regulates the HGs’ propensity to adopt farming practices from nearby farmers (see Methods).

We solved the model equations numerically to investigate the resulting gradient of Anatolian ancestry over the 3000 km transect. Published aDNA evidence suggests that post-Neolithic Europe exhibited significant Anatolian ancestry, exceeding 75% across all of Europe^39^. To evaluate which of our numerical solutions are consistent with this pattern, we assessed the average proportion of Anatolian ancestry after the transition to agricultural practices is complete—when the population of culturally defined HGs is nil. We based the diffusion constant, growth rate, and carrying capacities for both HGs and farmers on previous estimates from literature^21,43^. By altering the values of *f* and γ, we observed that the final Anatolian ancestry after the Neolithic expansion correlates most strongly with the ratio *C* = *f*/γ, rather than the individual parameters *f* or γ, even when *f* varies over four orders of magnitude (see Fig. 1). Moreover, an increase in the *f*/γ ratio strongly correlates with a decrease in Anatolian farming ancestry, falling below 50% when the ratio reaches 0.1—an outcome that is highly inconsistent with the predominant Anatolian ancestry proportion estimated from aDNA for post-Neolithic Europe^39^.

**Figure 1:**
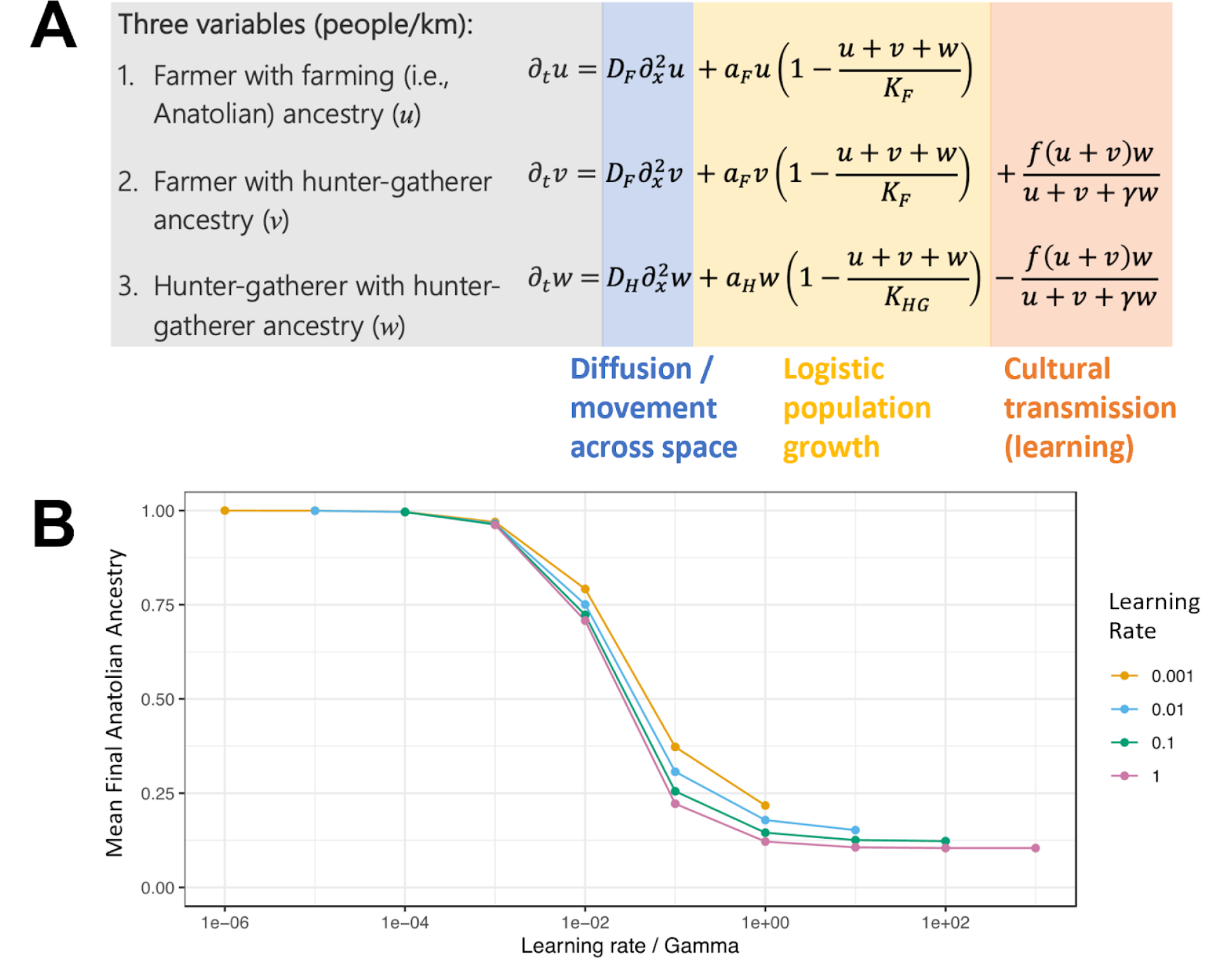
One-dimensional model (A) Our mathematical model utilizes a reaction-diffusion framework to simulate the spread of Anatolian ancestry across a one-dimensional European landscape initially populated by hunter-gatherers. It comprises a set of partial differential equations that describe the movement (diffusion) of individuals, their population growth (logistic), and the impact of cultural transmission on genetic ancestry. The model tracks three distinct groups: farmers with Anatolian ancestry, farmers with hunter-gatherer ancestry, and pure hunter-gatherers, each affected by diffusion constants, growth rates, and carrying capacities. The cultural transmission aspect is modeled through parameters that dictate the learning rate of new practices (f) and a bias toward adopting farming methods from either farmers or hunter-gatherers (γ). (B) Relationship between the ratio of the learning rate to the bias parameter (C = f/γ) and the proportion of Anatolian ancestry in the model, averaged across the landscape. The graph illustrates that the final proportion of Anatolian ancestry after the Neolithic expansion is predominantly determined by the ratio C, rather than the individual values of f or γ. Data points indicate model outcomes across a range of learning rates (f) and bias parameters (γ).

Given that the ratio of the learning rate to the bias parameter is the critical determinant in this model, we fixed γ to 1 (signifying unbiased learning) in subsequent analyses and focused solely on variations in the learning rate *f*. Thus, under unbiased learning, we find an upper limit to the learning rate of approximately 0.1 per generation (∼0.003 per year) for the proportion of Anatolian ancestry in our model to be consistent with the high levels observed in the aDNA data following the agricultural expansion.

Nonetheless, we acknowledge the model’s simplifications: an assumption of infinite population size, a single genetic locus without recombination, no age stratification or age-linked mortality, a unidimensional habitat, and exclusive mating within cultural groups. Subsequent sections will present simulations under more realistic conditions to verify the consistency of our findings and to refine our estimates of the extent of cultural transmission compatible with the archaeological record.

### Front speed is not informative about cultural transmission rate

To assess the validity of the conclusions under more realistic scenarios we turned to an agent-based model capable of simulating complex individual behavior (Fig. 2). This model, designed in SLiM (v4.0)^44^, considers unique individuals with sexually recombining genomes, age-based mortality, and local resource competition. Further, each individual moves and interacts on a customizable two-dimensional landscape that we utilize to simulate both simple and complex geography. Finally, the agent-based model has a cultural component in which HGs can learn and convert to agricultural lifestyles. This happens at a rate determined by the previously introduced learning rate parameter, *f*, and local farmer density (Fig. 2).

**Figure 2:**
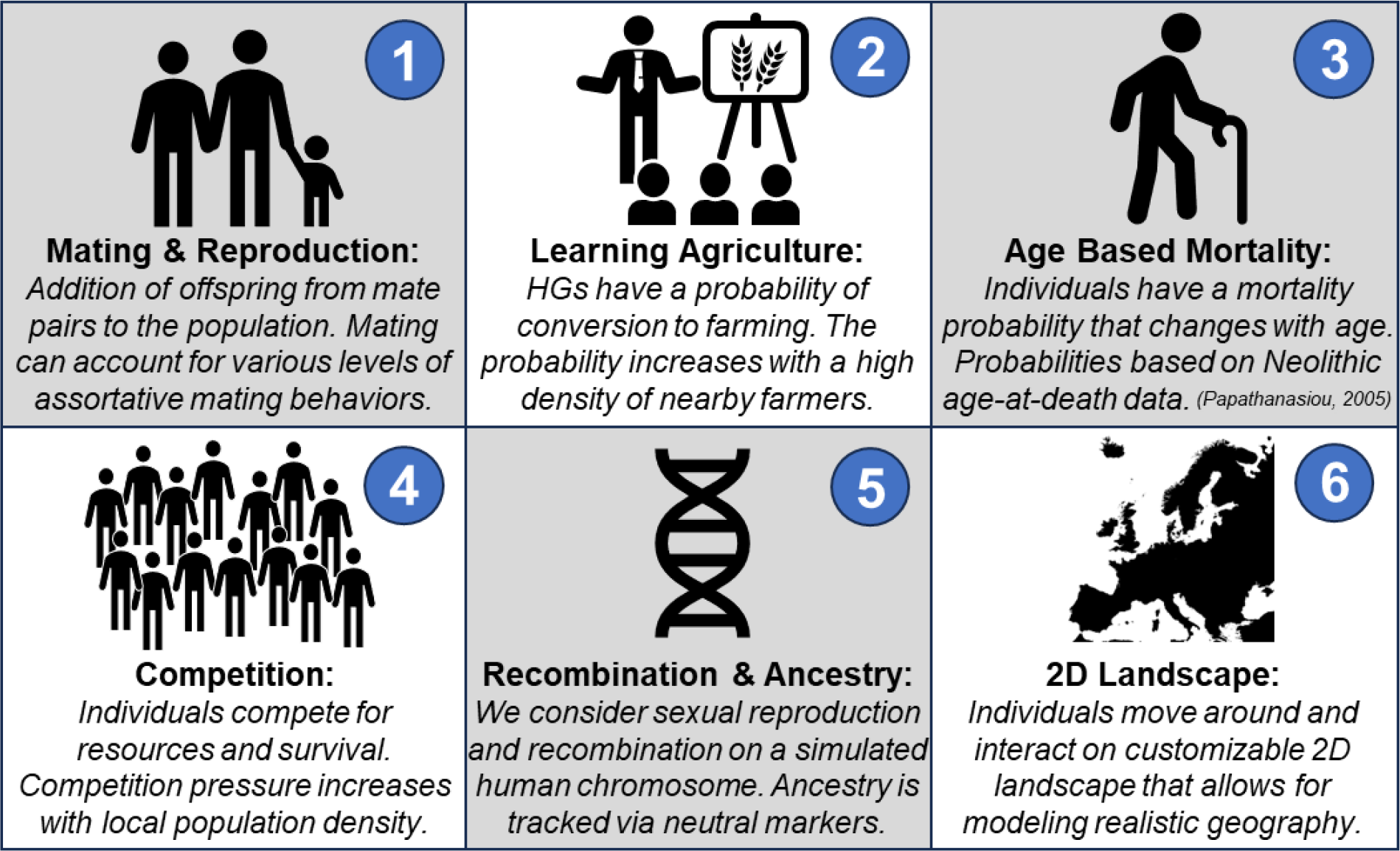
*Overview of the components in the agent-based simulation approach*. 1) Our model includes mating between individuals in the population under frequencies governed by assortative mating preferences and local abundance of cultural groups. Through reproduction, we simulate vertical (parent-offspring) cultural transmission. 2) We also consider hunter-gatherers learning agricultural behaviors via horizontal (peer-to-peer) cultural transmission. Learning occurs with a probability that is determined by a learning rate f and the local proportion of farmers surrounding a given hunter-gatherer. 3) Age-based mortality is implemented as a probabilistic mortality curve derived from an age-at-death study by Papathanasiou (2005)^34^. 4) Additionally, mortality is also governed by local competition for resources and considers different carrying capacities for farmers and hunter-gatherers. 5) Each individual has a simulated chromosome that is generated via simulated sexual recombination, allowing us to track ancestry proportions at an individual, and population, level. 6) Finally, the simulation is two-dimensional, with each individual having a location in space. Individuals exist and interact in a customizable virtual landscape.

As the simulation progresses, individuals move across the landscape and interact with each other (Fig. 3a). Each year, for every individual, a distance in the X direction (East-West) and a distance in the Y direction (North-South) is drawn from a normal distribution and added to the individual’s previous X, Y position. The standard deviations of the distributions (σ) allow us to control the distance individuals can travel within a year. We will primarily refer to σ as “step size”.

**Figure 3:**
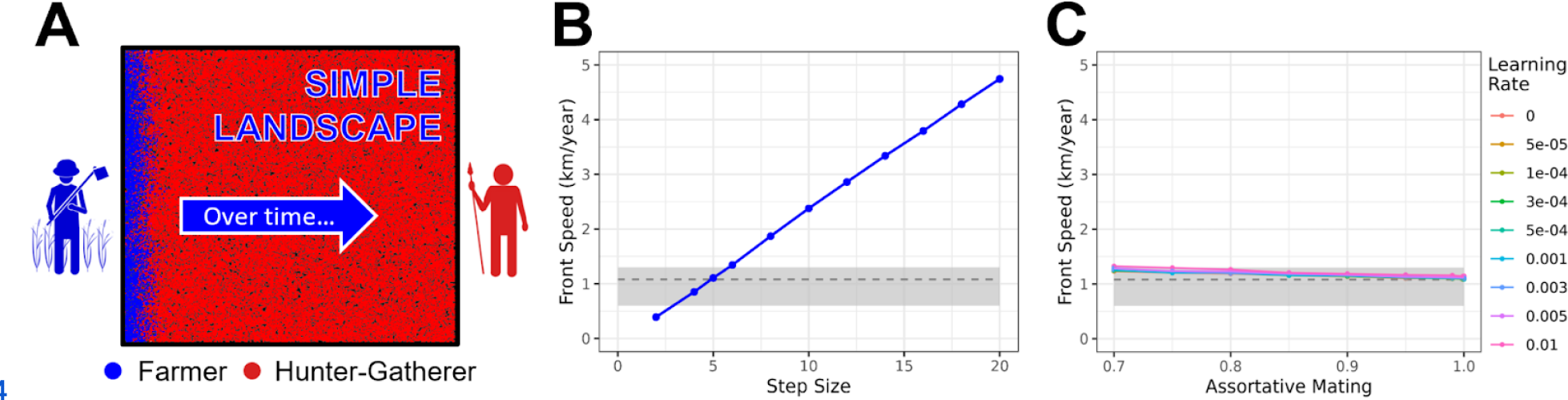
*Simple square landscape and the effect of step size, learning rate, and assortative mating on front speed* (A) Overview illustrating the progression of farming expansion on a simple square landscape from left to right. All front-speed calculations were done using the square landscape model depicted in panel A. Over time, the red hunter-gatherer individuals will be displaced (or converted) by blue farmers. (B) Effect of different σ values (referred to as step sizes) on front speed. For comparisons, the gray shaded box illustrates the front speed range of 0.6-1.3 km/yr estimated by Pinhasi et al. (2005)^22^ and the dashed line represents a front speed of 1.08 km/yr estimated by Ammerman and Cavalli-Sforza (1971)^20^. (C) Effect of assortative mating and different learning rates on front speed, given a fixed step size of 5 km. The gray shaded box again represents the front speed range of Pinhasi et al. (2005)^22^ and the dashed line the front speed estimate by Ammerman and Cavalli-Sforza (1971)^20^.

Similar to the diffusion constant in the mathematical model, the step size impacts the speed at which farmers diffuse across the landscape, and as a consequence, the speed of the cultural expansion (Fig. 3b). Based on estimates from the literature, we expected a front speed of approximately 1 km per year^20–22^ (gray box and dashed line in Figs. 3b & 3c).

To identify plausible σ values that are consistent with this front speed, we tested 11 values of σ between 2-20 km, in increments of 2 km, and calculated the front speed of each simulation under a fully demic model (see Methods). We found a σ of 5km to be the most plausible of the tested values (Fig. 3b) and used this step size as our standard throughout the rest of the simulations.

Next, we further explored the effect of assortative mating and learning on front speed (Fig. 3c). Using 5km as a fixed σ, we see that learning rates and assortative mating parameter values (within realistic ranges) have a negligible effect on the front speed. To affect front speed, very high rates of cultural transmission are required (Supplementary Fig. 1), however, these high rates almost completely inhibit the spread of Anatolian ancestry into Europe (Fig. 1) and thus are incompatible with empirical data^39^. We conclude that, although the front speed is informative about the step size parameter of the population, it does not provide fine-scale insight into the degree of cultural transmission. This is notable because it highlights the importance of co-analyzing ancestry and front speed in a combined model. In subsequent analyses, we explore the relationship between cultural transmission and genetic ancestry patterns and evaluate if ancestry patterns allow the estimation of cultural transmission parameters.

### Effect of cultural transmission on ancestry cline

We used our agent-based model to investigate the effect of assortative mating and learning on Anatolian ancestry proportions. Previous literature observed a cline of Anatolian ancestry with distance from the farming origin post-Neolithic Expansion^39^. In our simulations, we again assume that initially, all farmers are of pure Anatolian ancestry, and we explore how this ancestry spreads with the expansion of farming from left to right on a simple square landscape (Fig. 3a). We tested 16 learning rates in increments of 0.0003 between 0 and 0.005 (Supplementary Fig. 2) and restricted mating to be 100% assortative, i.e. by inhibiting matings between cultural farmers and HGs. All learning parameters result in a cline of Anatolian ancestry (estimated once all individuals are practicing farming) that decreases with increasing distance from the farming origin. As the learning rate increases, the slope of the ancestry cline also increases (Fig. 4a), such that learning rates larger than 0.003 result in less than 50% final Anatolian ancestry across most of the simulated landscape (Fig. 4a). This is consistent with our one-dimensional reaction-diffusion model that saw an equally strong reduction in final ancestry proportions for equivalent learning rates, further demonstrating that even low levels of learning can impede the spread of Anatolian ancestry.

**Figure 4:**
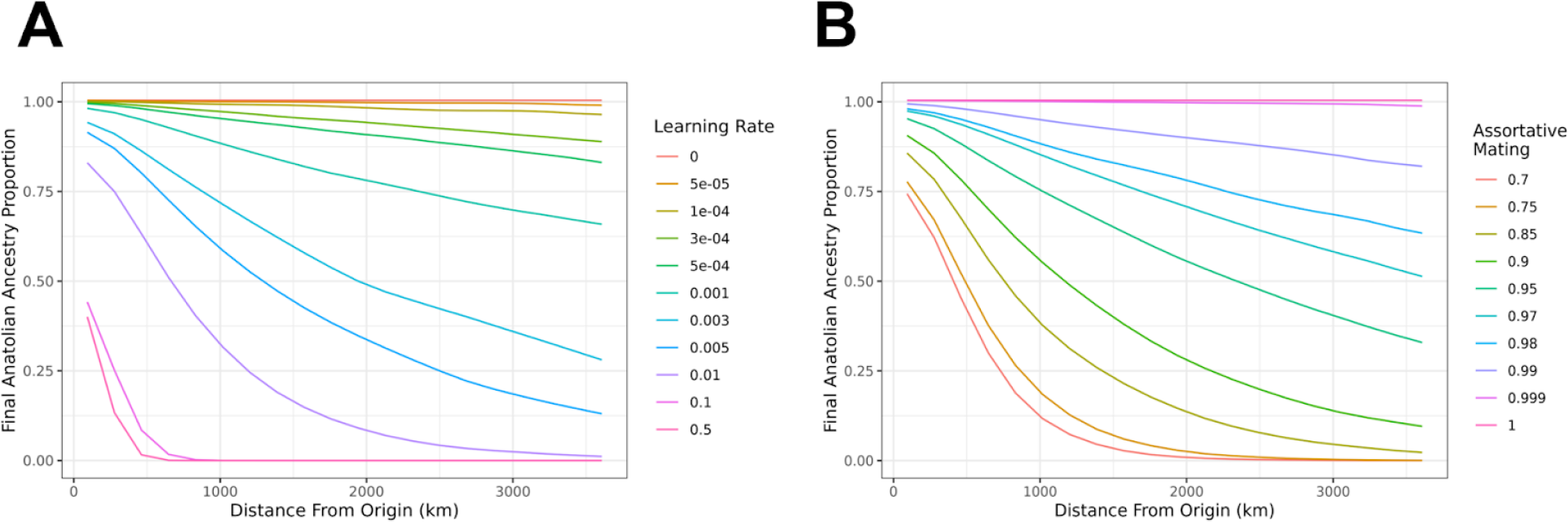
*Effect of cultural transmission on final Anatolian ancestry cline* Final Anatolian ancestry proportions plotted over distance from the origin of farming (km) upon the conclusion of simulations (when farming is ubiquitous across the landscape). (A) Simulations run with full assortative mating (100% assortative). Each line represents a different learning rate simulated. (B) Simulations run with zero learning but various assortative mating probabilities. Each line represents a different assortative mating probability.

To determine the sole effect of vertical cultural transmission on ancestry proportions, we fixed the learning rate to 0, i.e., restricting cultural transmission to cases where the offspring of a farmer and HG adopts farming. We then varied the assortative mating parameter, selecting 10 values between 0.7 and 1 (Supplementary Fig. 3). We found that low levels of non-assortative mating (i.e. HG with farmer) lead to similar ancestry clines (Fig. 4b) as the simulations with variable learning rates (Fig. 4a). This shows that both vertical and horizontal cultural transmission are analogous in their genetic effects. We further saw a rapid loss of Anatolian ancestry when we decreased the assortative mating rate, such that with assortative mating rates less than 90%, Anatolian ancestry quickly declined to zero toward the far end of the range. We observed similar decreasing Anatolian ancestry with increases in the learning rate, such that learning rates greater than 0.01 result in negligible Anatolian ancestry at the right end of the landscape.

In sum, both horizontal and vertical cultural transmission can generate spatial clines in Anatolian ancestry after the farming expansion, with higher rates of cultural transmission leading to steeper clines and lower final proportions. Given the proportion of Anatolian ancestry estimated from aDNA in post-Neolithic Europe is considerably high, we conclude that cultural transmission must have only played a limited role in the farming expansion.

### Estimating cultural transmission parameters from ancient DNA

Based on our aDNA estimations (see below) and published literature^39^, we know there to be a cline in Anatolian ancestry and not a complete replacement of HG ancestry after the farming expansion (see example ancestry cline Fig. 5a). This means some cultural transmission must have occurred, as a fully demic model would result in complete replacement. Because different learning rates lead to different clines of Anatolian ancestry (Fig. 4), we derived the learning rate that is most consistent with the observed European cline of Anatolian ancestry following the Neolithic Expansion^39^. To this end, we estimated Anatolian ancestry from a dataset of 911 aDNA samples from the Allen Ancient DNA Resource (AADR)^45^. We selected ancient samples that represent populations concurrent with and post the farming expansion but before the subsequent Steppe expansion^39,46^. Samples with the same geographic coordinates were binned together, resulting in 85 geographic sites, and Anatolian and Western Hunter-Gatherer (WHG) ancestry components were estimated for each site using the software qpAdm^47^ (see Methods). For accurate ancestry estimation, we only selected sites with more than 2 samples per site. All sites show Anatolian ancestry estimates close to or larger than 75% (Fig. 5b), except for three outlier sites with very low Anatolian ancestry (<50%). When plotting the estimated ancestry against direct distance from northwestern Anatolia (Ankara), a clear trend of decreasing ancestry with increasing distance can be observed (dashed gray line, Fig. 5b; slope = -6.200e-05, p-value = 7.09e-13). This is consistent with previous aDNA studies of the Neolithic^39^.

**Figure 5:**
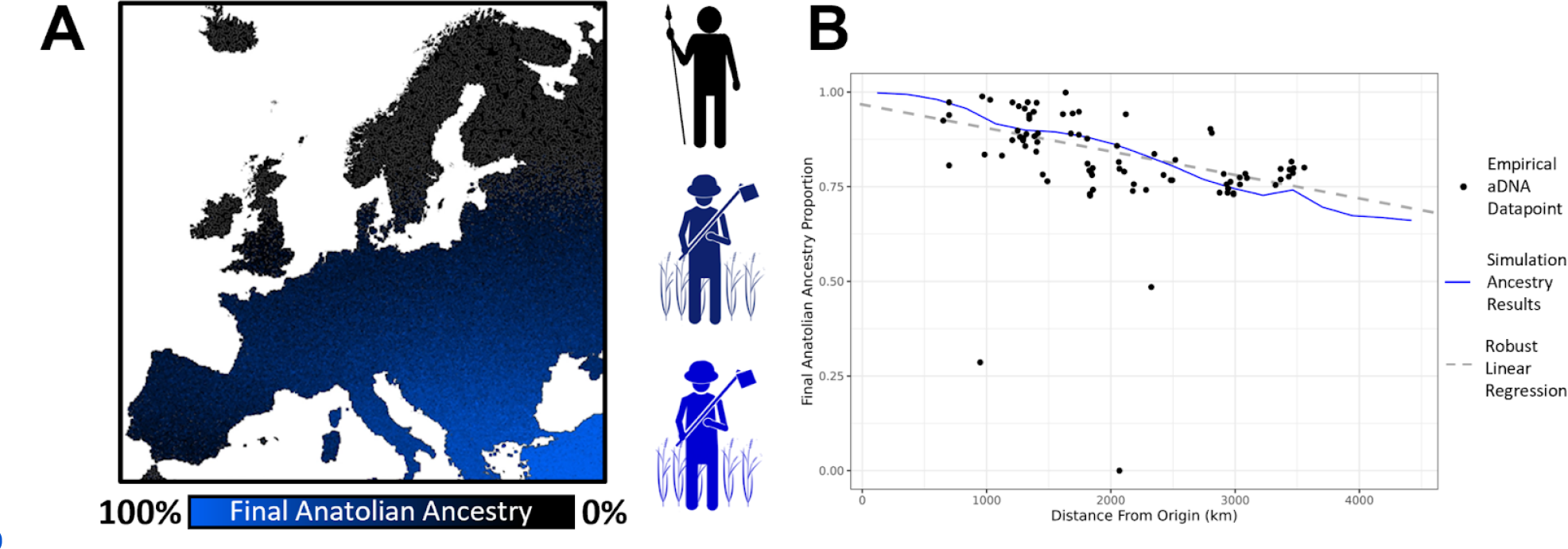
*Effect of cultural transmission on final Anatolian ancestry proportion on a European landscape* Final Anatolian ancestry proportion over distance from farming origin (km) plotted upon the conclusion of simulations when farming is ubiquitous across the landscape. (A) Figure showing a visual depiction of a hypothetical ancestry cline. Individuals on the map are colored by ancestry proportion. Blue represents high levels of Anatolian farming ancestry and black represents high levels of hunter-gatherer ancestry. Theoretical cline simulation run and depicted on an adapted map file from the European Environment Agency^48^. (B) Empirical aDNA ancestry estimates and best fitting simulation run. Black points show aDNA data points and their estimated ancestry proportions. The blue line shows the simulated ancestry results for the best-fitting simulation run (learning rate 0.00095). Simulations used the complex landscape map (Supplementary Fig. 6). The gray dashed line is a robust linear regression of the empirical aDNA data showing the significant negative cline of the Anatolian ancestry with increasing distance from the farming origin. The regression was performed using the ‘lmrob’ function from the R package ‘robustbase’ ^49^

Next, we used our “simple” square landscape model (Fig. 3a) to estimate a best-fitting learning parameter that is consistent with the ancestry cline observed from empirical data. Using the simulated ancestry rates shown in Fig. 4a, we computed the likelihood of each learning rate by taking into account site-specific point estimates and error rates of Anatolian ancestry extracted from the qpAdm analysis (see Methods). Likelihood values across samples were multiplied, assuming that estimation errors are independent across sites. A quadratic polynomial was then fit to the likelihood values to derive the maximum likelihood estimate and standard error of the learning parameter (Supplementary Fig. 4). This approach leads to a best-fitting learning rate of 0.000876 [0.000857, 0.000895] (Supplementary Fig. 4a). Using the same approach and the simulated clines in Fig. 4b, we also derived a best-fitting assortative mating parameter of 0.9836 [0.9832, 0.9839] (Supplementary Fig. 4b).

Our learning rate estimate assumes complete assortative mating such that only horizontal, but no vertical cultural transmission, takes place. Similarly, our assortative mating estimate assumes no learning and thus no horizontal transmission. Due to the equivalence of horizontal and vertical transmission in their effect on the ancestry cline (Fig. 4), this implies that our estimated learning rate can be considered an upper limit to the true learning rate in a combined model of both horizontal and vertical transmission, and vice versa for the estimated assortative mating rate (see Supplementary Fig. 5).

To enhance the realism of the simulated landscape, we subsequently generated data from a more detailed map, incorporating the outline of the European landscape and ensuring that individuals’ movement was confined within this outline (Supplementary Fig. 6). We based our simulations on the assumption of farming originating in Anatolia and tracked the ancestry cline as a distance from Anatolia following complete expansion of farming (see Methods). Finally, we repeated the same inference procedure as above and estimated a learning rate of 0.000951 [0.000931, 0.000971] (Supplementary Fig. 4c). Thus, the complex geography has only minimally impacted our parameter estimation when compared to the simple square landscape model. However, prior archaeological studies have indicated that the agricultural expansion was not uniform but might have spread faster along a coastal Mediterranean route compared to a northerly route through the Balkans and into Central Europe^37,50,51^. To evaluate how such a non-uniform expansion speed might have influenced our analysis, we ran simulations that decreased the step size parameter in the North-South direction to 4 km per year while raising the step size in the East-West direction to 6 km per year. Using the same maximum likelihood approach as before, we found a similar best-fitting learning rate of 0.000937 [0.000917, 0.000957] (Supplementary Fig. 4d).

In sum, we consistently estimate a yearly learning rate of less than 0.1%, suggesting that at the wavefront, effectively only 1 in 1,000 farmers converted an HG each year. This estimate is robust to modeling assumptions and one order of magnitude smaller than previous estimates of the learning rate based on the speed of the farming expansion alone^23,25^.

### Demic expansion without ancestry turnover?

We have shown that learning rates can vary substantially without affecting front speed (Fig. 3c), but that the observed expansion of Anatolian ancestry is only consistent with very low learning rates and almost complete assortative mating (Fig. 4 & Fig. 5b). Here, we more generally evaluate the contribution of cultural transmission to front speed in our simulations, and how this contribution relates to the turnover in ancestry. To this end, we evaluate the *cultural effect*, which is the contribution of cultural transmission to front speed relative to an otherwise demic model^23–25^:

Cultural effect (%) = (front speed - demic speed)/front speed ⨉ 100

We assessed the cultural effect for a wide range of learning rates under our simple landscape model (Fig. 6). We assumed complete assortative mating in these simulations but also explored a model of varying assortative mating rates (Supplementary Fig. 7). For our empirically estimated learning rate of approximately 0.001, the cultural effect is close to zero (0.5%), suggesting that this level of learning does not increase the front speed relative to a pure demic model. Thus, cultural transmission does not appear to have increased the expansion speed during the European Neolithic spread.

**Figure 6:**
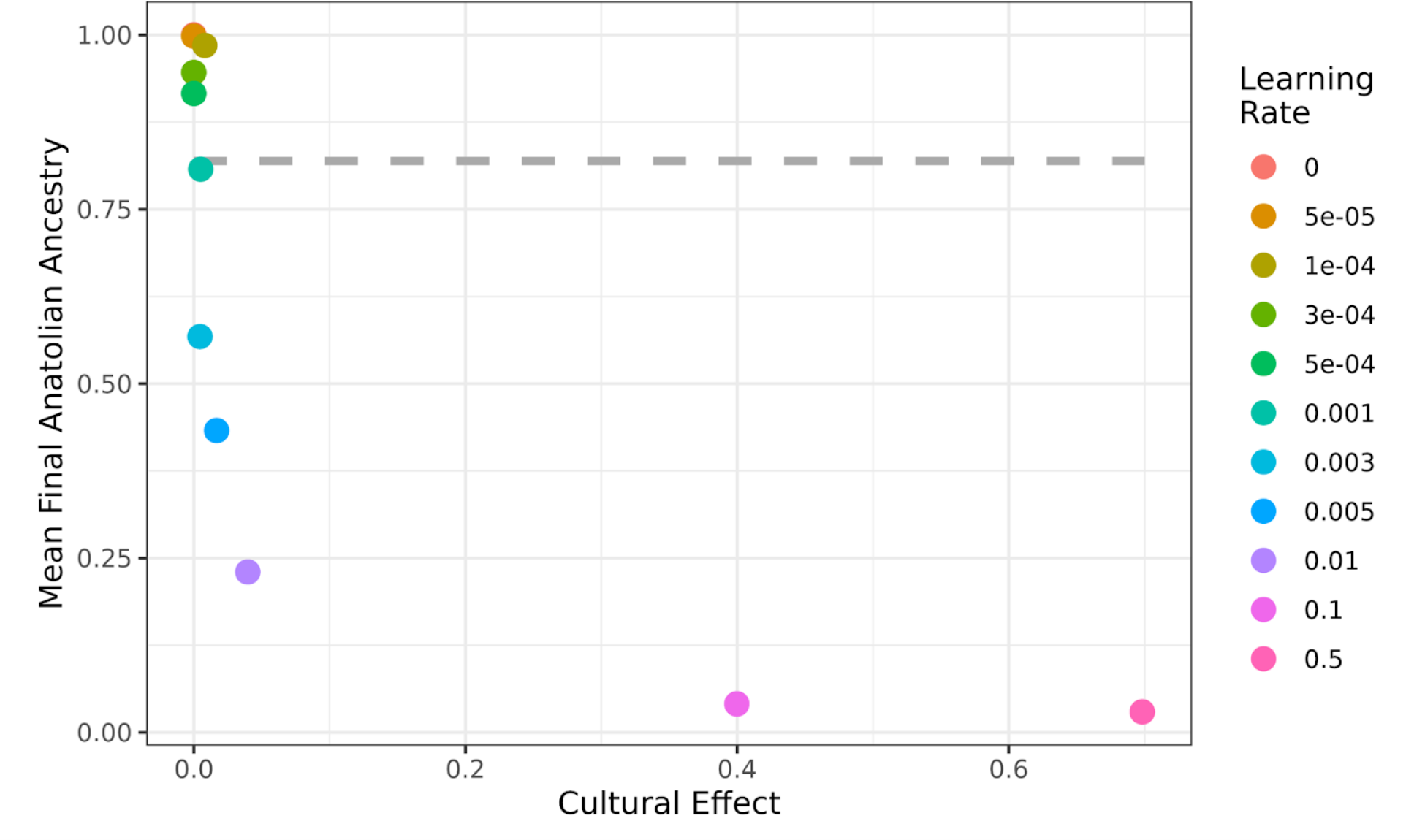
Mean final Anatolian ancestry proportion as a function of the cultural effect. Cultural effect is defined as the percent contribution of cultural transmission to the front speed, in relation to the baseline speed of a fully demic model. The figure shows the cultural effect under various learning rate parameters and the final Anatolian ancestry proportion upon the conclusion of the simulation when farming has become ubiquitous. Simulations were run with full assortative mating and various learning rates (right). For comparison to empirical data, the dashed gray line represents the mean proportion of Anatolian ancestry from our aDNA estimates.

However, when learning rate is further increased from this value, we see that for a large range of learning rates (i.e., *f* in [0.005, 0.1]) Anatolian ancestry does not spread across Europe despite the cultural effect being low. I.e., despite the model essentially being demic, it does not lead to a turnover in ancestry (Fig. 6). Only when *f* is larger than 0.1 does cultural transmission predominantly determine the front speed, indicated by a *cultural effect* larger than 50%. This is notable because a lack of ancestry turnover during a cultural expansion has been used previously as evidence for a cultural transmission model, e.g. regarding the spread of farming within the Near East^6,42^, or the expansion of the Bell Beaker culture from Iberia to Central Europe^52^. However, our results indicate that there is a wide range of cultural transmission rates that prevent a turnover of genetic ancestry, but are small enough to lead to a predominantly demic diffusion of culture (*cultural effect* < 50%), i.e. geographic ancestry patterns do not necessarily change during a demic expansion of culture.

## Discussion

While archaeological and genetic research has described the impacts of the Neolithic transition from their respective disciplines, our interdisciplinary approach enables us to understand cultural and behavioral mechanisms invisible to the material evidence. Our mathematical and simulation-based modeling of the Neolithic expansion, for the first time, simultaneously predicts archaeological and genetic patterns linked to one of the most significant transformations in human history.

Our study focuses on the extent to which cultural transmission facilitated the widespread adoption of agricultural practices during the Neolithic Revolution in Europe. Using reaction-diffusion modeling and agent-based simulations, we show that the expansion of agriculture across Europe was predominantly driven by a continuous process of farmer migration and demographic replacement, with only a minimal contribution of cultural transmission of agricultural practices. Specifically, we estimate a remarkably low cultural transmission rate— at the wavefront, only 1 in every 1,000 farmers annually converted a Hunter-Gatherer (HG) to agriculture. This rate is significantly lower than inferences from earlier studies based solely on the speed of the expansion, which estimated that 3.3% to 8.3% of farmers each year were converting HGs, assuming a 30 year generation time^23^.

Another metric for quantifying the relative contribution of different mechanisms in the spread of farming is the “cultural effect”. Cultural effect is the extent to which cultural transmission impacts the front speed within a demic-cultural framework^23–25^. We estimate that the cultural effect is negligibly small (0.5%), whereas prior studies have reported estimates from 21% to 40%^23,25^. However, these prior estimates fail to consider the distinct ancestry patterns that would result from such high levels of cultural transmission. Moreover, previous approaches are often underpinned by untestable assumptions about the movement of people and how culture was transmitted. For example, they utilize the distance between spousal birthplaces to calculate dispersal ranges^23,24,53^, or studies of modern missionary trips to determine how readily HGs convert to farming^23^, assuming that these quantities are also applicable to prehistoric populations. Our model does not rely on such assumptions, instead fitting parameters to empirical observations of front speed and ancestry.

However, our model does not consider the possibility of individuals reverting from farming back to hunting and gathering, maintaining that once an individual adopts farming, their descendants are invariably farmers. This assumption, while grounded in ethnographic evidence^54,55^ and consistent with previous models^25^, may oversimplify the transmission of cultural practices. We tested our model under alternative conditions, i.e., where offspring could adopt either lifestyle of their two parents, yet found similar ancestry clines, indicating a limited impact of this assumption on our results (Supplementary Fig. 8). Overall, our modeling implies that mating between cultural groups must have been remarkably rare (<2% non-assortative mating; Supplementary Fig. 4b). Notably, this is corroborated by aDNA studies which do not find any farmer-associated ancestry in hunter-gatherers that overlapped in time with neighboring Neolithic farmers^56,57^.

Our model further assumes a uniform European landscape, thereby potentially glossing over critical geographic factors that could influence local ancestry distribution patterns. For example, previous studies suggested a deceleration in the spread of farming in northern latitudes, attributed to several factors^28–30,51^. Climate has been posited as a significant influence, shaping agricultural practices, interactions between distinct groups, and crop performance^51^. Additionally, the presence of denser HG populations in these regions has been proposed as a competitive barrier to the northward agricultural spread^28–30^. This deceleration of the farming expansion was linked to higher retention of HG ancestry in northern latitudes, suggesting that the slower rate of expansion allowed for greater integration or coexistence between incoming farmers and indigenous HG communities^51^. In our model’s assessment of non-uniform expansion, conducting additional simulations with a decreased step size of farmers indeed demonstrated that a slower expansion speed leads to greater HG ancestry retention in the population following the expansion (Supplementary Fig. 9). This may explain why in northern regions where the farming expansion was presumably slower, e.g. due to the southwest Asian crop package underperforming in the colder and wetter climate, higher levels of HG ancestry are found^51^. Importantly, estimated learning rates remained consistently low in our model, irrespective of whether we assumed uniform dispersal or a faster latitudinal expansion (Supplementary Fig. 4d).

Our study primarily examines the agricultural expansion in Europe. In the global context, Neolithic expansions exhibited considerable variability in their replacement of native HG ancestry, challenging the notion that agricultural spread always was accompanied by significant genetic turnover^58^. In fact, the agricultural expansion within southwest Asia, before its spread into Europe, is not associated with ancestry turnover, which has led to the conclusion that this initial expansion was propelled more by the dissemination of ideas and farming technology than by the movement of people^6^. However, our simulations suggest complexities beyond conventional classifications into cultural vs. demic diffusion based on ancestry turnover, revealing scenarios where agricultural expansion—despite being predominantly demic—does not lead to the anticipated widespread dissemination of farmer ancestry. Instead, significant portions of HG ancestry can be maintained even under a predominantly demic expansion model.

Consequently, our findings prompt a reevaluation of other significant cultural shifts, such as the spread of the Bell Beaker phenomenon around 4,800-4,000 years ago. Similar to the farming expansion in southwest Asia, the Bell Beaker cultural phenomenon initially spread throughout Western Europe without affecting genetic ancestry— individuals associated with Bell Beaker sites have large regional variations in ancestry. However, the later expansion of the Beaker complex to Britain led to a massive replacement of almost all of Britain’s gene pool. Again, this was interpreted as a cultural expansion that was initially mediated by cultural diffusion^52^. However, our modeling suggests that a wide range of cultural transmission rates can prevent a turnover of genetic ancestry— but are small enough to lead to a mainly demic diffusion of culture (i.e. with a cultural effect < 50%; Fig. 6). Thus, even the initial expansion of the Beaker complex is, in principle, consistent with a predominantly demic mechanism. In sum, our insights call for a reconsideration of how we understand cultural spreads and their impact on genetic ancestry patterns, potentially revising narratives around major prehistoric and historic cultural expansions.

In conclusion, our research offers an estimate of the cultural contribution to the Neolithic agricultural expansion in Europe, suggesting that the front speed was not significantly increased by cultural transmission. Through the application of diverse modeling approaches—including reaction-diffusion and agent-based models, alongside simulations of complex geography and varied European expansion axes—we reveal that previous estimates of a cultural effect of 21-40%^23,25^ are likely overstated. Our analysis, incorporating both front speed and empirical ancestry data from ancient DNA, suggests that only a minute fraction of farmers might have been actively involved in cultural interactions with HGs. More generally, our modeling stresses the possibility of demic expansion without complete ancestry turnover, offering a refined perspective on historical expansions like the Bell Beaker phenomenon in Europe. Our interdisciplinary approach allows us to more accurately dissect the dynamics of the Neolithic expansion, emphasizing the need for integrated methodologies and modeling approaches in historical analysis.

## Methods

### One-dimensional equation model

We set up a reaction-diffusion model, based on partial differential equations, that incorporates spatial diffusion of individuals, cultural transmission of farming practices, and genetic ancestry at a single locus. Under this model, we assumed a 3000 km long one-dimensional (1D) habitat along which the density of three types of individuals is modeled: cultural farmers with Anatolian ancestry, cultural farmers with HG ancestry, and cultural HGs with HG ancestry. As an initial condition, we assume that the farmers of Anatolian descent are constrained to approximately 200 km at one end of the range, and the HG group is constrained to the remaining 2,800 km of the range, with about a twenty-fold difference in population density^21^. Our mathematical model assumes a diffusion component, a logistic population growth component, and a cultural transmission component (see formulas in Fig. 1a). The cultural transmission model was originally developed by Fort 2012^23^, but without accounting for the ancestry of individuals. The cultural transmission component of our model decreases the HG density and to the same extent increases the “farmers with HG ancestry” density as a function of two parameters, the learning rate *f* and the bias parameter *γ*. The learning rate *f* is a product of the number of teachers an HG contacts during their lifetime, and the probability that an HG converts to farming after contact with a farmer. The bias parameter *γ* modifies the preference of HGs to learn from a farmer rather than from another HG^23^. The diffusion constant *D* was chosen such that the expansion speed of farming under purely demic parameters is the empirically observed 1 km per year, given a growth rate ɑ of 3% per year^21,59^. We assumed that both farmers and HGs have the same diffusion constant and growth rate. In accordance with the initial condition, we assumed that the carrying capacity of farmers was twenty times larger than that of HGs^21^. We set up the partial differential equations model in Mathematica 13.3^60^ and numerically solved it over a time span of 150 generations (i.e., ∼4500 years when assuming a generation time of 30 years) using the Mathematica function *NDSolve*. The boundary conditions of the numerical solution were set up such that at positions 0 km and 3000 km (the left and right boundary of the 1D landscape), the first derivative of the three population densities with respect to the space coordinate was set to zero across all time points.

### Two-dimensional agent-based simulation model

Using the insights from the basic 1D reaction-diffusion model, we then developed a more realistic two-dimensional (2D) agent-based model, written in the Eidos language^61^ designed for the SLiM simulation framework^44^ (v4.0). In these simulations, each individual exists on a simulated 2D landscape, has a diploid genome, and a cultural identity (farmer or HG). We adopt annual timesteps and model overlapping generations, meaning that individuals persist across multiple yearly intervals until they pass away. Upon initialization of the simulation (year one), the ancestry assigned to the individuals’ chromosomes is based on their initial cultural identity (i.e., farmers have pure Anatolian ancestry and HGs have pure native HG ancestry). In each subsequent year, the ancestry proportion of new offspring is determined via sexual recombination of the individual’s parental genomes. The cultural identity of each individual is independent of their genetic ancestry and can change from HG to farmer through means of cultural exchange as the simulation progresses.

### Genome simulation

To determine the ancestry proportion of each individual, neutral marker mutations are initialized every megabase along the 247Mb simulated chromosome (human chromosome 1) of all farmers. These markers “tag” Anatolian farming ancestry along the chromosome of each individual. Accordingly, the proportion of Anatolian ancestry is calculated by counting the number of markers and dividing by the total number of marker locations. The genomes of new individuals are generated via simulated sexual recombination of the two parents as part of the built-in SLiM reproduction function^44^, assuming a recombination rate of 1 cM/Mb.

### Mortality and competition

Mortality in our agent-based model is governed by two main factors: age-dependent mortality and density-dependent competition. Our age-dependent mortality curve is based on osteological age-at-death data collected by Papathanasiou (2005)^34^. We generated an age-specific ’equilibrium’ mortality rate that corresponds to the age-at-death distribution observed by Papathanasiou^34^ (Supplementary Table 1), i.e. in equilibrium, this mortality rate leads to the age-at-death distribution observed by Papathanasiou^34^. Using these ’equilibrium’ mortality rates for each age, we computed an equilibrium fertility rate of 0.1, for each mature individual over the age of 11. I.e., this fertility rate compensates for yearly deaths, which allows the population size to stay constant over time. The ages of individuals at the initiation of the simulation were sampled from the equilibrium age distribution.

To model density-dependent competition and convergence of the population size towards a carrying capacity *K*, we implemented a density-dependent scaling of the ’equilibrium’ mortality curve. To this end, all age-dependent mortality rates were scaled down in regions with population density below *K*, and scaled up in regions with population density above *K*. The local population pressure is calculated as the ratio of the number of neighbors within a radius of 30 km surrounding each individual, relative to the expected local *K* within this area. We derive each individual’s ’experienced’ mortality rate by multiplying its ’equilibrium’ mortality rate by the local population pressure, which allows the population to grow until it hits the carrying capacity *K*. Each cultural group has a unique *K*— *K_F_* for farmers, and *K_HG_* for HGs, which allows farming to support higher population densities.

We set the *K_HG_* to 0.064 individuals per square kilometer. This is following the modeling approach by Currat and Excoffier (2005)^43^, who selected this value based on two previous papers exploring palaeo-demographics^62,63^. For the farmer carrying capacity (*K_F_*) we used a value of 1.28 individuals per square km. We calculated this number based on estimates by a model by Ammerman and Cavalli-Sforza (1984). In this model, the *K_F_*is expected to be 20 times larger than *K* ^21^.

In our model, density-dependent competition takes into account all individuals, independent of cultural identity. As a result, HG populations gradually decline over time as the population pressure from closeby farming populations becomes too large for their smaller *K_HG_*. This is based on the idea that farmers compete with HGs for land and resources, which was also incorporated in previous mathematical models of the farming expansion^28–30^. To reduce the runtime and memory requirements of the agent-based simulation we downscaled the number of individuals by a factor of five. Thus, we divided all carrying capacities mentioned above, as well as the starting population density, by 5 for all agent-based simulation runs.

### Reproduction and assortative mating

Each year, each mature individual randomly chooses a mate within its cultural group (assortative mating), or independent of cultural group (non-assortative mating), governed by an assortative mating probability parameter set before runtime. Additionally, mate selection only occurs between mature individuals (age > 11) within a radius of 10 km. When a suitable mate is found, then the number of offspring is sampled from a Poisson distribution with a fertility rate of 0.1. This rate is calculated based on the age-specific mortality curve, such that population size is maintained when the population is at carrying capacity (i.e., deaths are compensated by births; see above) (Supplementary Table 1). If no suitable mate is found (e.g. if an HG assortatively mates, but is not surrounded by any other HG within a radius of 10 km), then no offspring is produced by that individual in that year, except it is selected as a mate by another individual. The location of offspring is chosen as one of the parents location, simulating maternal care.

If a HG and a farmer reproduce, their offspring is assigned to be a farmer. This assumption is supported by cultural observations of the assimilation of individuals into farming groups^54,55^, and the lack of farming ancestry in the genomes of HGs in ancient DNA studies^56,57^. However, we have validated the robustness to this assumption with a model where offspring of farmer-HG matings randomly choose their subsistence strategy (Supplementary Fig. 8).

### Learning and cultural transmission

Each year, every HG individual has the opportunity to learn agriculture and transition to being a farmer. The individual probability of transitioning to farming takes into account the local proportion of farmers surrounding the HG individual in a radius of 10 km (i.e., the more farmers there are in the vicinity, the higher the probability of learning; when no farmers are nearby, learning will not take place). Thus, for each HG in the simulation at a given year, the transitioning to farming is a random event with probability calculated as the product of the local proportion of farmers and a simulation-wide learning rate *f*. This model is consistent with the cultural transmission model of Fort 2012^23^. The product of the fixed number of teachers encountered each year and the probability of learning farming from a single farmer contact, in their model, equals to our learning rate *f*. It can be interpreted as the ’number of HGs converted per farmer’ when the local proportion of farmers is close to zero (i.e. at the tip of the wave-front of the farming expansion), or alternatively as the proportion of HGs that gets converted to farming when farming is the dominant local subsistence strategy. Further note that our rates are yearly rates, whereas Fort 2012 models cultural transmission on a per-generation basis^23^.

### Simulation landscape

The landscape map in our model is represented by a black-and-white image file that is uploaded to the SLiM^44^ software. The black pixels represent inhabitable land, and the white represents uninhabitable regions (i.e. oceans). Within our 2D spatial model, each individual has a positive X and Y coordinate that represents their location on the inhabitable landscape. The 2D position of the individuals is compared to the location on the landscape map, allowing us to achieve geographic boundaries for expansion.

We utilized this feature to run simulations on both a simple all-black pixel plane where all regions of a 3,700 x 3,700 km landscape are inhabitable, as well as a map outlining the geography of Europe. The latter map was adapted from the European Environmental Agency’s (EEA) Elevation map of Europe^48^, available at the following URL: (https://www.eea.europa.eu/ds_resolveuid/558D91E1-3DB0-4639-9F70-2012CC4453A5). Using Adobe Photoshop (CC 2022)^64^, we made the landscape image file dichromatic by selecting the land areas and filling them with black pixels, whereas we changed the remaining pixels (i.e., oceans) to white. To facilitate water travel via ocean, we added water crossings to the map using the directional maps from Fort (2015) as reference^24^ (Supplementary Fig 6).

### Movement

Each year, for every individual, a distance in the X direction (East-West) and a distance in the Y direction (North-South) is drawn from normal distributions with standard deviations σ*_X_* and σ*_Y_*, which are then added to the individual’s previous X, Y position. These standard deviations allow us to control the distance individuals can travel within a year. Whenever the coordinates of the newly sampled position are out of bounds (i.e., off the map or on a white pixel of the landscape), this position is rejected and a new position is sampled until a legitimate position is found. For most simulations, we assume isotropy, i.e. σ*_X_* and σ*_Y_*are the same. However, we explore simulations with σ*_X_* > σ*_Y_* to investigate the impact of a favored East/West movement direction, e.g. reflecting the faster expansion along the Mediterranean route^37,50,51^.

### Front speed calculation

Our simple square landscape was used to investigate how front speed, i.e. the speed with which farming expands from the left to the right side of the square, depends on step size and learning rate parameters. Upon initialization, we filled up the landscape with individuals in accordance with the HG carrying capacity (*K_HG_*). Individuals with X-coordinates between 0 and 74 km were designated as original farmers, while the rest of the individuals on the map were initialized as HGs. We then tracked the expansion of farming each year by calculating the proportion of farmers within twenty 185 km wide bins on the X-axis. The extent of farming expansion was defined as the X coordinate of the bin at which the farmer proportion had just reached 50%. The speed of farming expansion was then calculated by the slope of a linear regression of the extent of farming against time.

### Empirical ancestry estimates

We used qpAdm from the ADMIXTOOLS2^47^ package with parameter ‘allsnps=TRUE’ to obtain empirical estimations of Neolithic Anatolian and European Western Hunter-Gatherer ancestry contributions to early European farmers from ancient DNA (Allen Ancient DNA Resource (AADR) v52.2)^45^. We formed the European HG source population by grouping the LaBrana1 (sample ID I0585)^7^, Loschbour.DG^41^, & KO1 (sample ID I1507)^7^, and formed the Anatolian Neolithic population from 26 samples from the archaeological sites of Barçın Höyük^7^ and Boncuklu Tarla^65^. We supplied the following groups as fixed right-group populations in qpAdm: Mbuti.DG^66^, Papuan.DG^67^, Han.DG^66^, Karitiana.DG^66^, Ethiopia_4500BP.SG^68^, Italy_North_Villabruna_HG^69^, Russia_Ust_Ishim.DG^70^, Czech_Vestonice^69^, Russia_MA1_HG.SG^71^, and Israel_Natufian^6^. We grouped the target populations based on the labels provided in the AADR V52.2 annotation data column Group ID and filtered any singleton groups and groups with a mean age between 5000 and 8000 BP. We report scaled qpAdm admixture weight estimates whereby to keep the values between zero and one we multiplied each estimate by 1 / the sum of both weights.

### Fitting simulations to empirical ancestry data

To fit paramters of our simulations to the qpAdm^47^ results, we selected aDNA data sampled after the farming expansion but before the Steppe expansion^39,46^ (4,500-8,000 ybp) and compared the empirical Anatolian ancestry estimates to the simulated ancestry clines (i.e., the Anatolian ancestry proportion as a function of the distance to Anatolia) under different learning and assortative mating rates. Thus, for each aDNA sample in our study, we derived an observed Anatolian ancestry proportion using the empirical estimates from qpAdm^47^, and we derived an expected Anatolian ancestry proportion, assuming the same distance to Anatolia, from our agent-based simulations. To this end, we took the straight-line distance of each ancient sample from Ankara and used our simulation data to interpolate an expected Anatolian ancestry proportion with the same straight-line distance on the simulated map. We took into account uncertainty in the estimation of the empirical ancestry proportion by assuming a normally distributed measurement error with standard deviation set to the sample-specific standard error output of qpAdm^47^. For each simulated learning rate, the observed and expected Anatolian ancestry proportions were used to calculate a log-likelihood of each ancient sample, which was then summed up across all samples. The simulation-wide log-likelihood was then plotted against each respective learning rate. We fit a quadratic function to the log-likelihood values to derive a maximum likelihood estimate of the learning rate and a standard error as the square root of the negative inverse of the second derivative (Supplementary Fig. 4).

Note that to derive the expected Anatolian ancestry proportion from our simulations, we had to bin simulated individuals into different distance bins to compute their average ancestry proportion as a function of distance. For the simple square landscape, we binned all individuals into 20 equal-sized bins along the X-axis, since farming progresses along the X-axis from left to right in this model. On the simulated European map, where farming starts in Anatolia and then expands into Europe, we created bins based on the straight-line distance of each individual from the origin in the southeast corner of the map, representing Anatolia. We then sampled 10,000 individuals randomly throughout the map and binned them according to this distance into 20 bins with 262km width, and for each bin computed the average Anatolian ancestry.

## Author Contributions

T.M.L. and C.D.H. conceptualized and designed the study, T.M.L. developed and analyzed the 2D agent-based simulations, C.D.H. developed the 1D mathematical model, M.P.W conducted aDNA ancestry estimations, T.M.L. and C.D.H. prepared the manuscript, T.M.L., C.D.H. and M.P.W. revised and edited the manuscript, C.D.H. supervised the study. All authors read and approved the final manuscript.

## Funding

Research reported in this publication was supported by the National Institutes of Health Grant R35GM146886, by the Pennsylvania State University, and by the National Institutes of Health T32 GM102057 Computation, Bioinformatics, and Statistics (CBIOS) Training Program Grant. The content is solely the responsibility of the authors and does not necessarily represent the official views of the National Institutes of Health.

## Code Availability Statement

The Edios code for the SLiM agent-based model used in this study, a file explaining the code for the SLiM agent-based model— as well as landscape files for the model, the R data analysis scripts, and the Mathematica code for the 1D model, can all be found in the following repository: https://github.com/TroyLaPolice/European-Neolithic-Expansion

## Competing Interest Statement

The authors declare no competing interest.

## Notes

https://github.com/TroyLaPolice/European-Neolithic-Expansion

### Competing Interest Statement

The authors have declared no competing interest.

