## Supplementary Information / Extended Data for "The European Neolithic Expansion: A Model Revealing Intense Assortative Mating and Restricted Cultural Transmission"

***Supplementary Table 1:*** *Age-based mortality probability values generated from Neolithic osteological age at death data*[1].

| Age | Mortality Probability at this Age | Probability of Survival to this Age |
|---|---|---|
| 0 | 0.046336 | 0.953664396 |
| 1 | 0.046336 | 0.90947578 |
| 2 | 0.046336 | 0.86733467 |
| 3 | 0.046336 | 0.827146194 |
| 4 | 0.046336 | 0.788819876 |
| 5 | 0.056409 | 0.744323656 |
| 6 | 0.056409 | 0.702337405 |
| 7 | 0.056409 | 0.66271954 |
| 8 | 0.056409 | 0.625336462 |
| 9 | 0.056409 | 0.590062112 |
| 10 | 0.022 | 0.577081023 |
| 11 | 0.022 | 0.564385513 |
| 12 | 0.022 | 0.551969298 |
| 13 | 0.022 | 0.539826233 |
| 14 | 0.022 | 0.527950311 |
| 15 | 0.035354 | 0.509285058 |
| 16 | 0.035354 | 0.4912797 |
| 17 | 0.035354 | 0.473910906 |
| 18 | 0.035354 | 0.457156173 |
| 19 | 0.035354 | 0.440993789 |
| 20 | 0.056801 | 0.415945066 |
| 21 | 0.056801 | 0.392319126 |
| 22 | 0.056801 | 0.370035154 |
| 23 | 0.056801 | 0.349016926 |
| 24 | 0.056801 | 0.329192547 |
| 25 | 0.069353 | 0.306362189 |
| 26 | 0.069353 | 0.285115177 |
| 27 | 0.069353 | 0.265341699 |
| 28 | 0.069353 | 0.246939564 |
| 29 | 0.069353 | 0.229813665 |
| 30 | 0.090704 | 0.208968618 |
| 31 | 0.090704 | 0.190014303 |
| 32 | 0.090704 | 0.172779222 |
| 33 | 0.090704 | 0.157107435 |
| 34 | 0.090704 | 0.142857143 |

| | | |
|---|---|---|
| 35 | 0.122008 | 0.125427464 |
| 36 | 0.122008 | 0.110124341 |
| 37 | 0.122008 | 0.096688318 |
| 38 | 0.122008 | 0.084891594 |
| 39 | 0.122008 | 0.074534161 |
| 40 | 0.160622 | 0.06256236 |
| 41 | 0.160622 | 0.052513489 |
| 42 | 0.160622 | 0.044078685 |
| 43 | 0.160622 | 0.036998692 |
| 44 | 0.160622 | 0.031055901 |
| 45 | 0.167447 | 0.02585569 |
| 46 | 0.167447 | 0.021526237 |
| 47 | 0.167447 | 0.017921738 |
| 48 | 0.167447 | 0.0149208 |
| 49 | 0.167447 | 0.01242236 |
| 50 + | 1.0 | 0.0 |

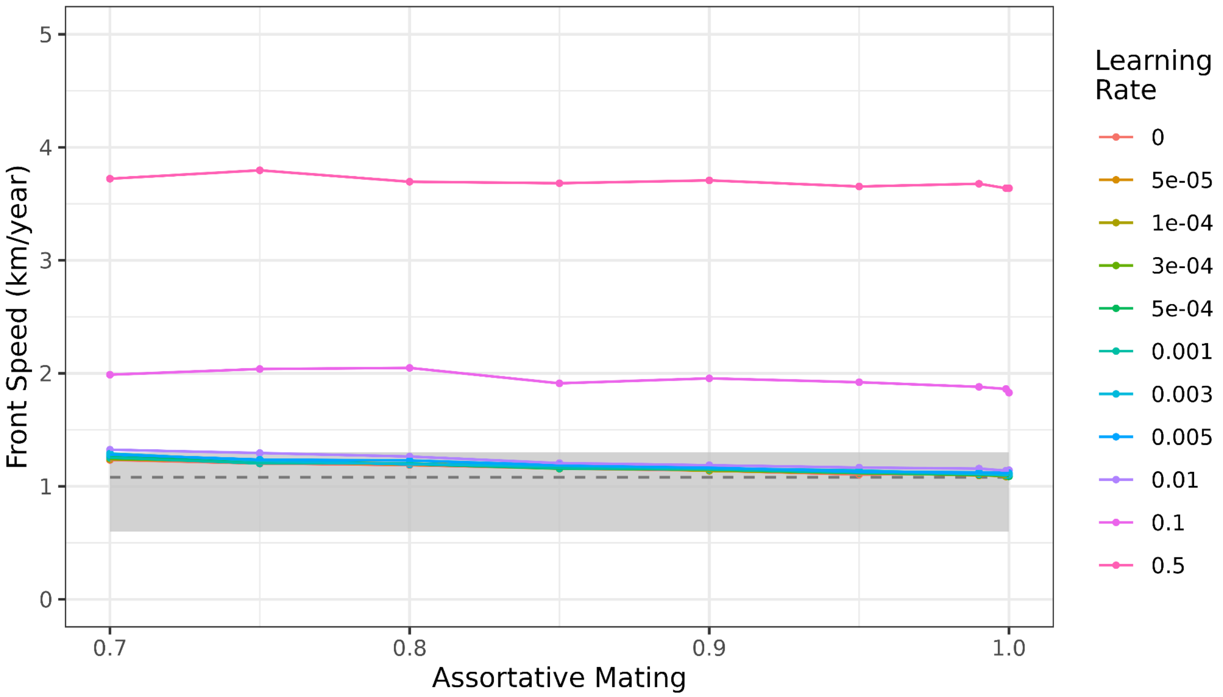


***Supplementary Figure 1: Effect of learning rates and assortative mating on front speed***

*Effect of assortative mating and different learning rates on front speed, given a fixed step size of 5 km. For comparisons, the gray shaded box illustrates the front speed range of 0.6-1.3 km/yr estimated by Pinhasi et al. (2005)[2] and the dashed line represents front speed of 1.08 km/yr predicted by Ammerman and Cavalli-Sforza (1971)[3]. This supplementary plot shows additional very high learning rates not shown in Fig. 3c (0.1, 0.5) that are required to affect front speed.*

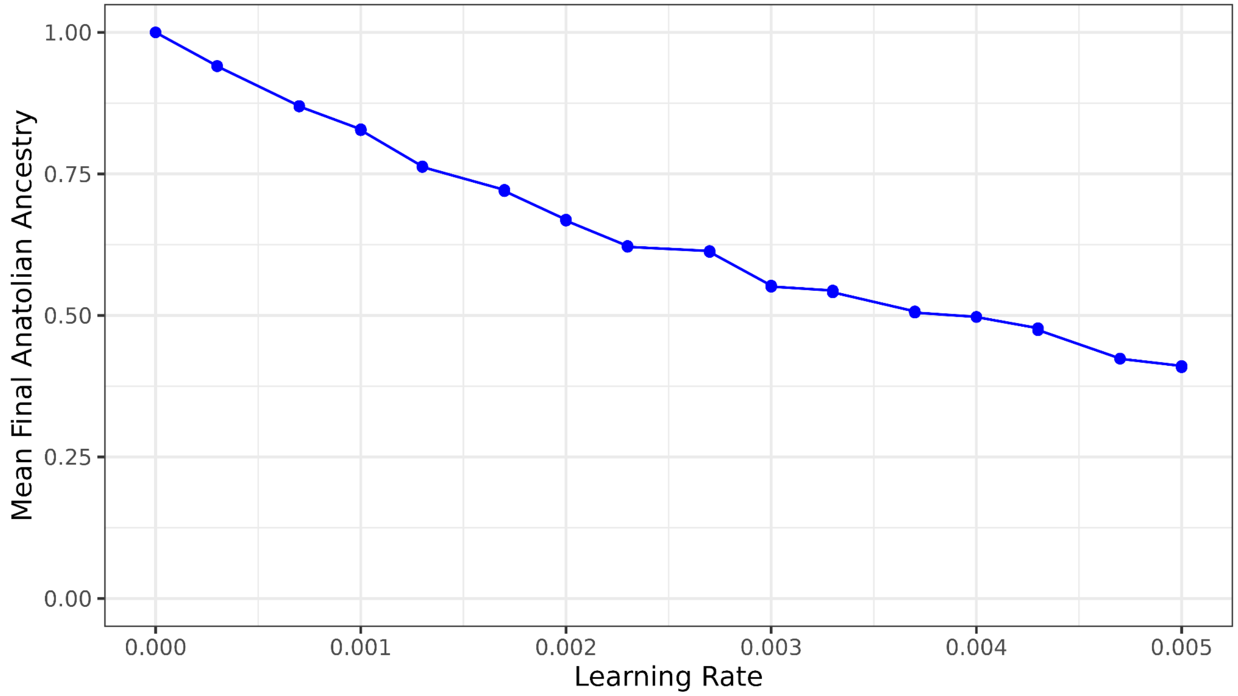


***Supplementary Figure 2: Effect of learning rates on remaining Anatolian ancestry***
*16 simulated learning rates used for fitting empirical data and their effect on the proportion of Anatolian ancestry in the population following the expansion, when farming has become ubiquitous. Figure shows the mean proportion of Anatolian ancestry in the population across the entire landscape. All 16 runs assume fully assortative mating. All runs used the simple square landscape.*

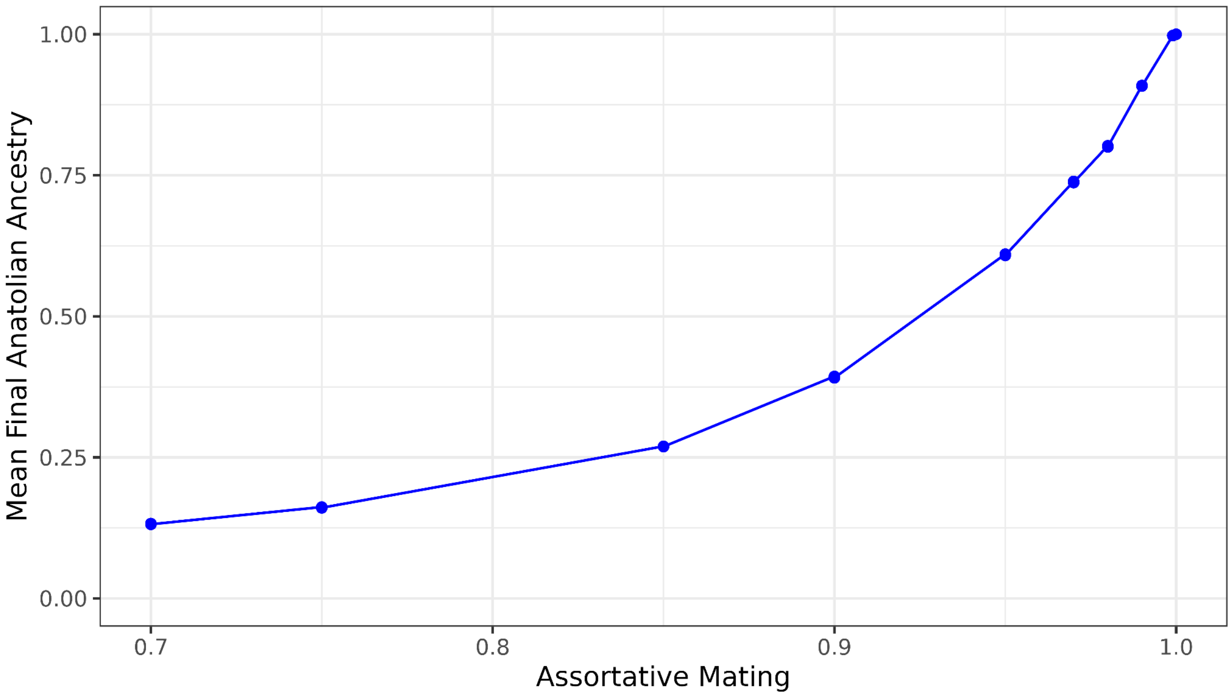


***Supplementary Figure 3: Effect of assortative mating rates on remaining Anatolian ancestry***
*10 simulated assortative mating rates used for fitting empirical data and their effect on the proportion of Anatolian ancestry in the population following the expansion, when farming has become ubiquitous. Figure shows the mean proportion of Anatolian ancestry in the population across the entire landscape. All 10 runs assume zero peer-to-peer learning with only parent-to-child vertical cultural transmission. All runs used the simple square landscape.*

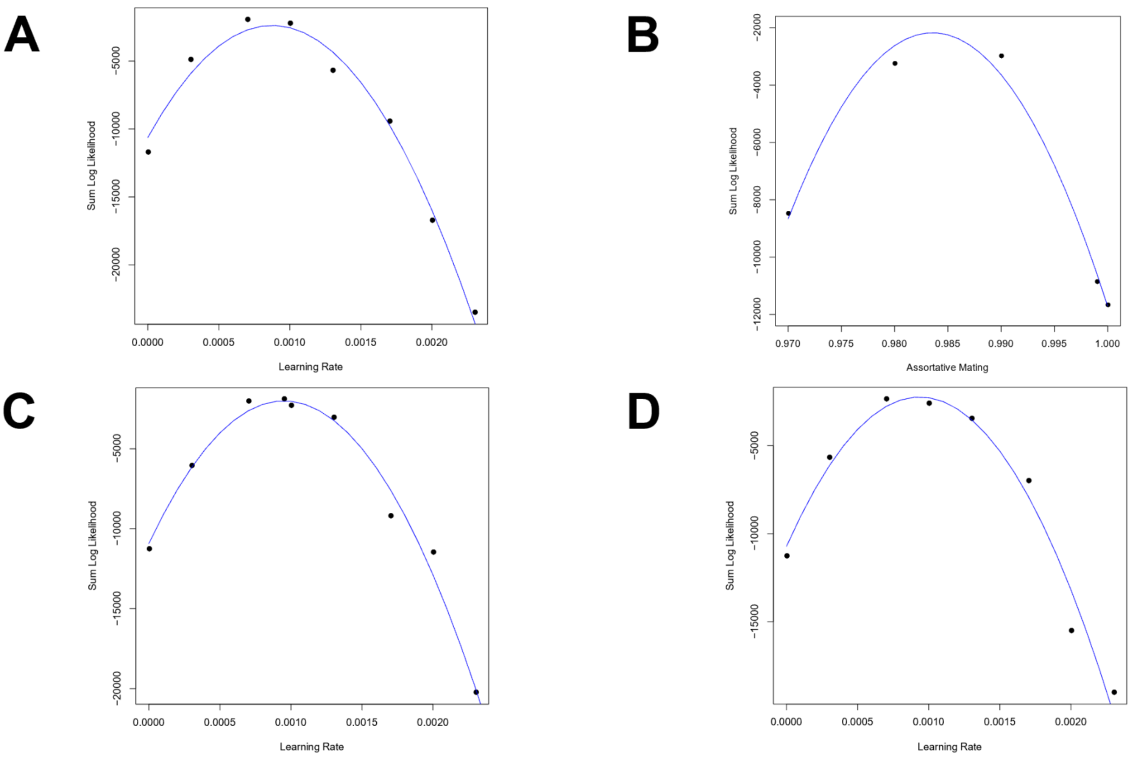


***Supplementary Figure 4: Quadratic polynomial fit to the log-likelihood values to determine the maximum likelihood estimates for best-fitting parameters***

*(A) Learning rate estimates for simple landscape map. (B) Assortative mating estimates for simple landscape map. (C) Learning rate estimates for complex landscape map. (D) Learning rate estimates for complex landscape map with farmer movement biased along a Mediterranean expansion route. In all cases, we selected 5-9 log-likelihood points close to the optimum to enable a more accurate local fit of the quadratic approximation.*

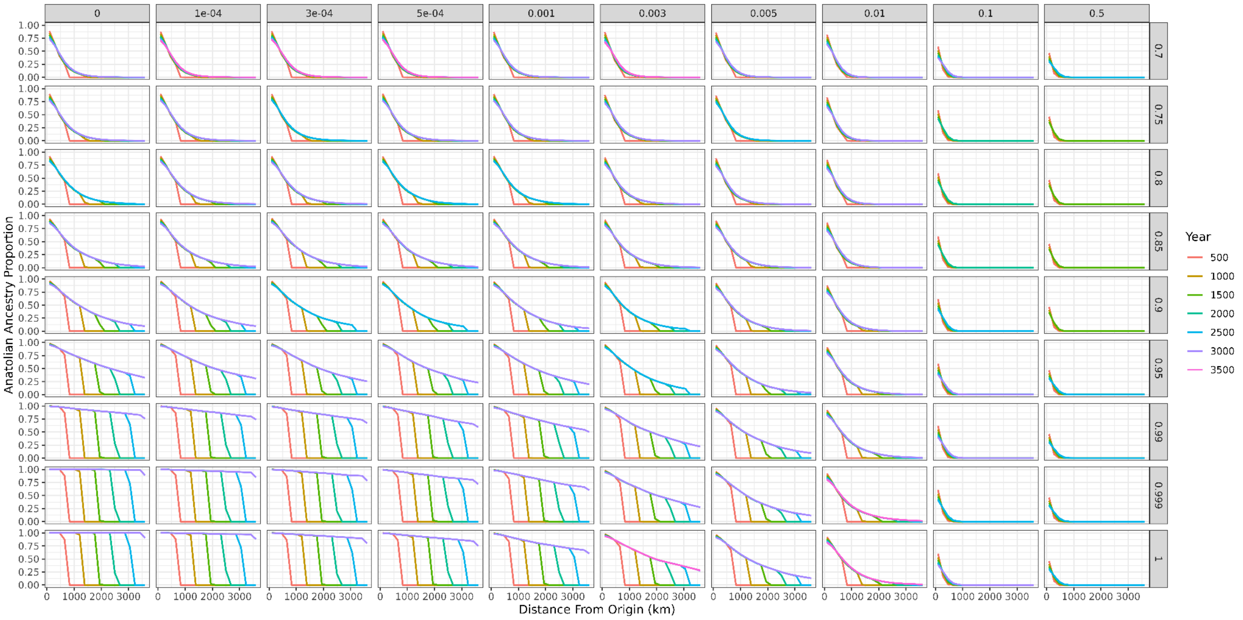


***Supplementary Figure 5: Interaction of learning and assortative mating parameters***

*Y-axes show the Anatolian ancestry proportion in the population through the course of the simulation plotted over distance from origin (km). Lines show the progression of the expansion over time with different colored lines plotted in 500 year increments. Plot faceted on the x-axis by learning rate and on the y-axis by assortative mating parameters tested.*

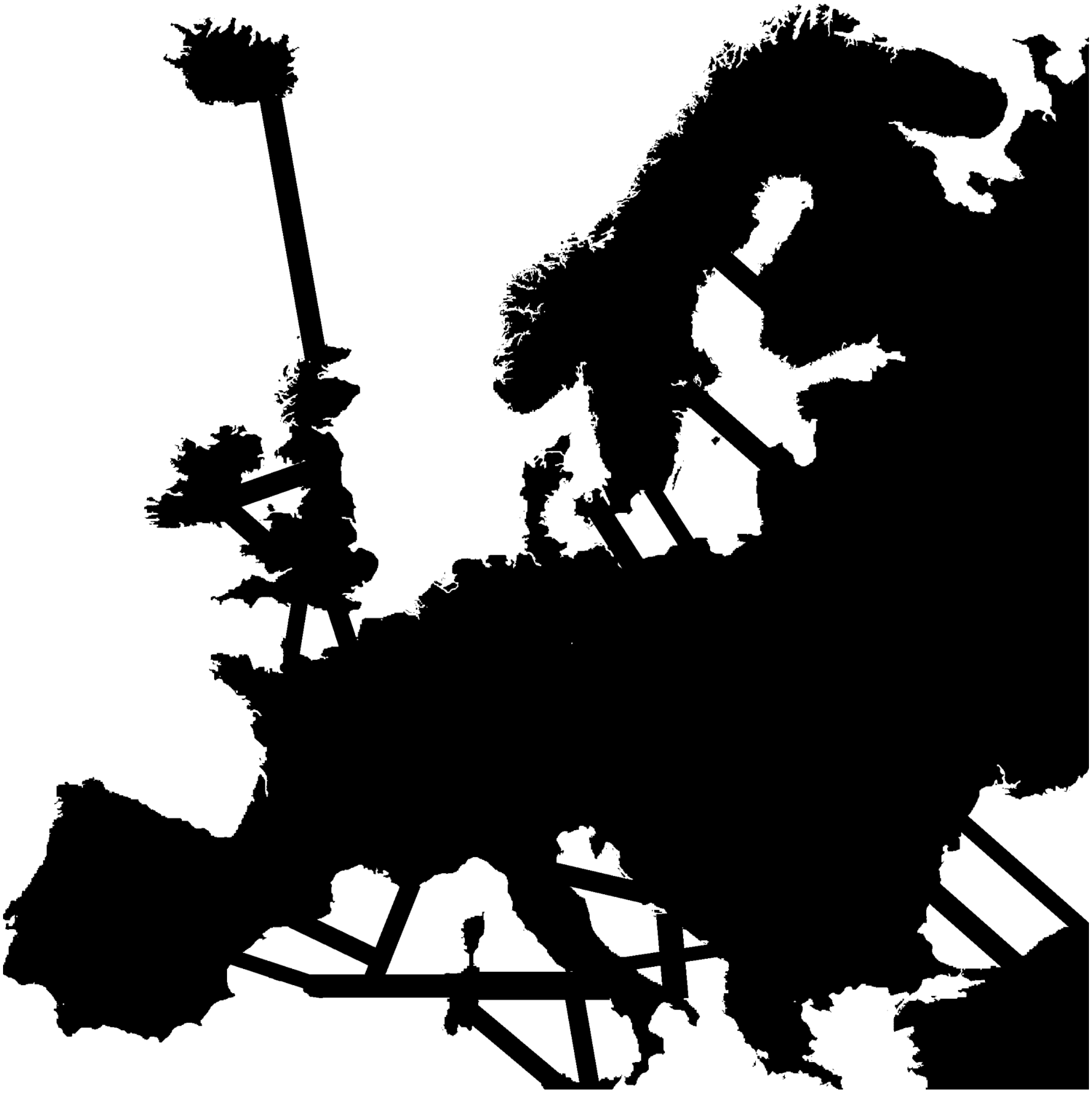

***Supplementary Figure 6: 2D landscape map for complex geography simulations***
*Complex map runs were conducted on this map file. The visible "land bridges" seen in the Mediterranean (and elsewhere) facilitated water crossings that would have taken place via boat, but in the simulation were larger than the possible yearly step size for individuals. Map file adapted from the European Environmental Agency's (EEA) Elevation map of Europe[4], available at the following URL:*
(https://www.eea.europa.eu/ds_resolveuid/558D91E1-3DB0-4639-9F70-2012CC4453A5).

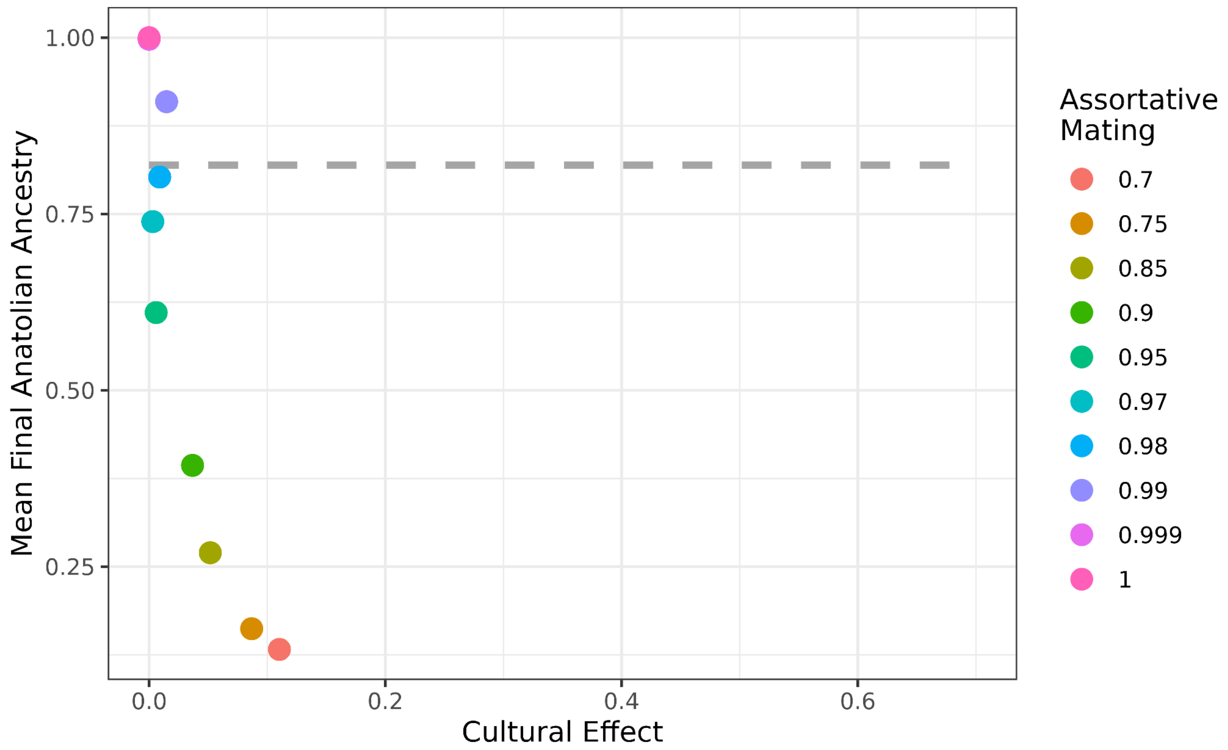


***Supplementary Figure 7: Mean final Anatolian ancestry in the population as a function of the cultural effect given various assortative mating rates***

*Cultural effect describes the percent contribution of cultural transmission to the front speed in addition to a baseline speed of a fully demic model. This figure shows the cultural effect under various assortative mating parameters and the mean Anatolian ancestry proportion across the simulated population upon the conclusion of the simulation when farming has become ubiquitous. Simulations were run with no peer-to-peer learning and only parent-to-child vertical cultural transmission via various assortative mating rates (right). For comparison to empirical data, the dashed gray line represents the mean proportion of Anatolian ancestry from our aDNA estimates.*

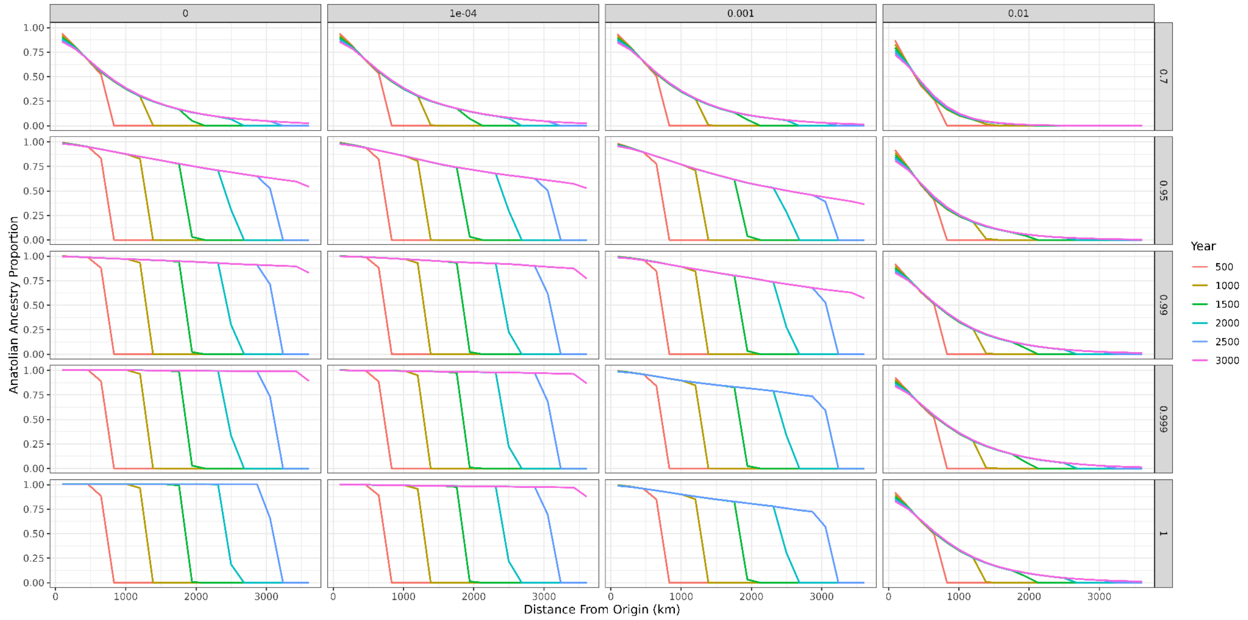


***Supplementary Figure 8: Interaction of learning and assortative mating parameters without the assumption that children of farmers only become farmers***

*These plots are results from simulations where the offspring of a farmer and a HG has a 50/50 chance of becoming either a HG or a farmer. Other simulations in the study assume that any non-assortative matings between HGs and farmers always result in a farming offspring. These plots do not assume farming offspring. The panels show Anatolian ancestry proportion in the population, over the course of the simulation, plotted over distance from origin (km). Lines show the progression of the expansion over time with different colored lines plotted in 500 year increments. Plot faceted on the x-axis by learning rate and on the y-axis by assortative mating parameters tested.*

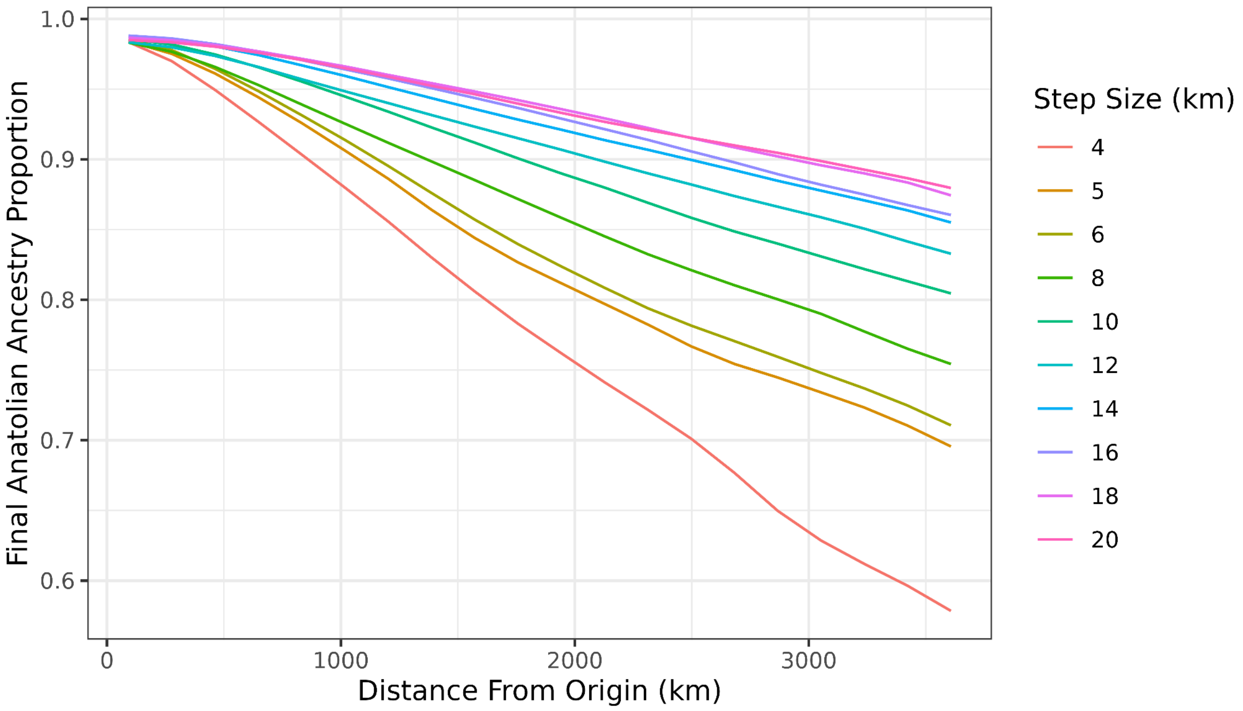


***Supplementary Figure 9: Effect of different farmer step sizes on Anatolian farmer ancestry proportion over distance from farming origin***

*Final Anatolian ancestry proportions plotted over distance from origin (km) upon the conclusion of simulations (when farming is ubiquitous across the landscape). All simulations are run with full assortative mating and a learning rate of 0.001. Each colored line represents a different step size parameter.*
